# Scorepochs: A Computer-Aided Scoring Tool for Resting-State M/EEG Epochs

**DOI:** 10.1101/2020.05.26.116434

**Authors:** Matteo Fraschini, Simone Maurizio La Cava, Giuseppe Rodriguez, Andrea Vitale, Matteo Demuru

## Abstract

M/EEG resting-state analysis often requires the definition of the epoch length and the criteria to select which epochs to include in the subsequent steps. However, the effects of epoch selection remain scarcely investigated and the procedure used to (visually) inspect, label, and remove bad epochs is often not documented, thus hindering the reproducibility of the reported results. In this study, we present Scorepochs, a simple and freely available tool for the automatic scoring of resting-state M/EEG epochs that aims to provide an objective method to aid M/EEG experts during epoch selection procedure. We tested our approach on a freely available EEG dataset containing recordings from 109 subjects using the BCI2000 64-channel system.

## 1. Introduction

Task-free resting-state recording from M/EEG (magneto/electroencephalogram) represents one of the most used experimental paradigms to investigate the baseline level of brain activity in healthy subjects and patients [1]. However, the resting-state condition is an elusive concept which is influenced by different states of vigilance that are usually out of control from the experimenter [2]. Generally, the first steps performed during an M/EEG resting-state analysis consist of (i) segment the raw/filtered EEG traces into a set of non-overlapping epochs and (ii) select a number of artifacts-free epochs to be used during the subsequent steps of the pipeline. These steps require the definition of the epoch length and the criteria to select which epochs to include in the successive analysis. The effects of the epoch length have been previously investigated [3], while the effects of epoch selection, induced by the inter-observer variability and unclear criteria used for this task, remain scarcely investigated. Epoch selection is performed at individual level (independently for each subject) and usually is conducted by one or more experts. Some kind of procedure to detect and mitigate EEG artifacts may be applied before this step. However, the precise procedure used to (visually) inspect, label, and remove bad epochs is often not documented [4], thus hindering the reproducibility of the reported results. Most importantly, especially in the case the selection procedure is performed by different experts, it would be of relevance to assure that homogeneous criteria were used. In this context, most of the studies using resting-state paradigms make assumptions on stationarity of EEG signals and perform averaging of individual features (extracted at subject-level) to make inferences at group-level. In short, this means that it is assumed a strong within-subject stability of M/EEG features and that these individual characteristics may be consistent within a group. Subjective visual scoring and/or inter-observer variability poses a possible threat to the validity of these assumptions, although some studies have reported that the subjective influence may lead to minimal changes when a sufficient number of epochs are selected [5,6], nevertheless, it is still far to be clear how to quantify this sufficient number of epochs. In this context, it will assume a very important role the possibility to develop some kind of semi-automated analysis with the aim to help clinicians and researchers during these very crucial steps. Few recent studies used computer-assisted tools to allow interpreting EEG background patterns [7,8] but, probably, as suggested by van Diessen et. al. [4], these methods were not applied to large scale also because of the inherent complexity or limited transparency. On the other hand, other studies have suggested the use of automatic artifacts suppression [9] that would indirectly help to limit the uncertainty induced by inter-observer variability. Most of these approaches are based on independent component analysis (ICA) [10], which however requires a great amount of EEG data to obtain an acceptable decomposition (a minimum of 20 time points per channel^2) [11]. In this study, we present Scorepochs, a simple and freely available tool for the automatic scoring of resting-state M/EEG epochs that aims to provide an objective method to aid M/EEG experts during epoch selection. Our approach, which works at subject-level, provides a score for each epoch within a single M/EEG recording, with the attempt to make this crucial procedure less ambiguous, more objective, and reproducible. Scorepochs is based on the whole power spectrum of the EEG, does not require any specific assumption on the underlying frequency content and may thus keep all the relevant spectral information contained in the unfiltered raw signal [12].

## 2. Methods

The proposed method is based on a very simple and fast algorithm which takes as input (*i*) a set of M/EEG recordings and (*ii*) the length of the desired epoch. After that, the algorithm provides as output a score for each single M/EEG epoch. A schematic representation of the proposed method is depicted in Figure 1. Furthermore, all the scripts used to perform the analysis are freely available for MATLAB (https://github.com/Scorepochs-tools/scorepochs_mat) and for Python (https://github.com/Scorepochs-tools/scorepochs_py). For each subject, each epoch, and each channel, the algorithm computes the power spectral density (PSD) via the Welch method into a specific range of frequencies (see Figure 1, panels A and B). At channel-level, a similarity score, computed by using the Spearman correlation coefficient, is evaluated between the PSD extracted from all the epochs, thus providing a correlation matrix with *number of epochs x number of epochs* as dimension (see Figure 1, panel C). A first average is now computed over the rows (columns) of the symmetric matrix to obtain a *score vector* with length equal to the *number of epochs*, where the entries represent the mean similarity score of the corresponding epoch (see Figure 1, panel D). Computing the *score vector* for each channel, and then averaging the *score vectors* across channels, it is possible to obtain a final score for each epoch (see Figure 1, panel E). Finally, for each subject, the score can be sorted in descending order allowing to select the suggested epochs to be included in the subsequent steps of the analysis.

**Figure 1.**
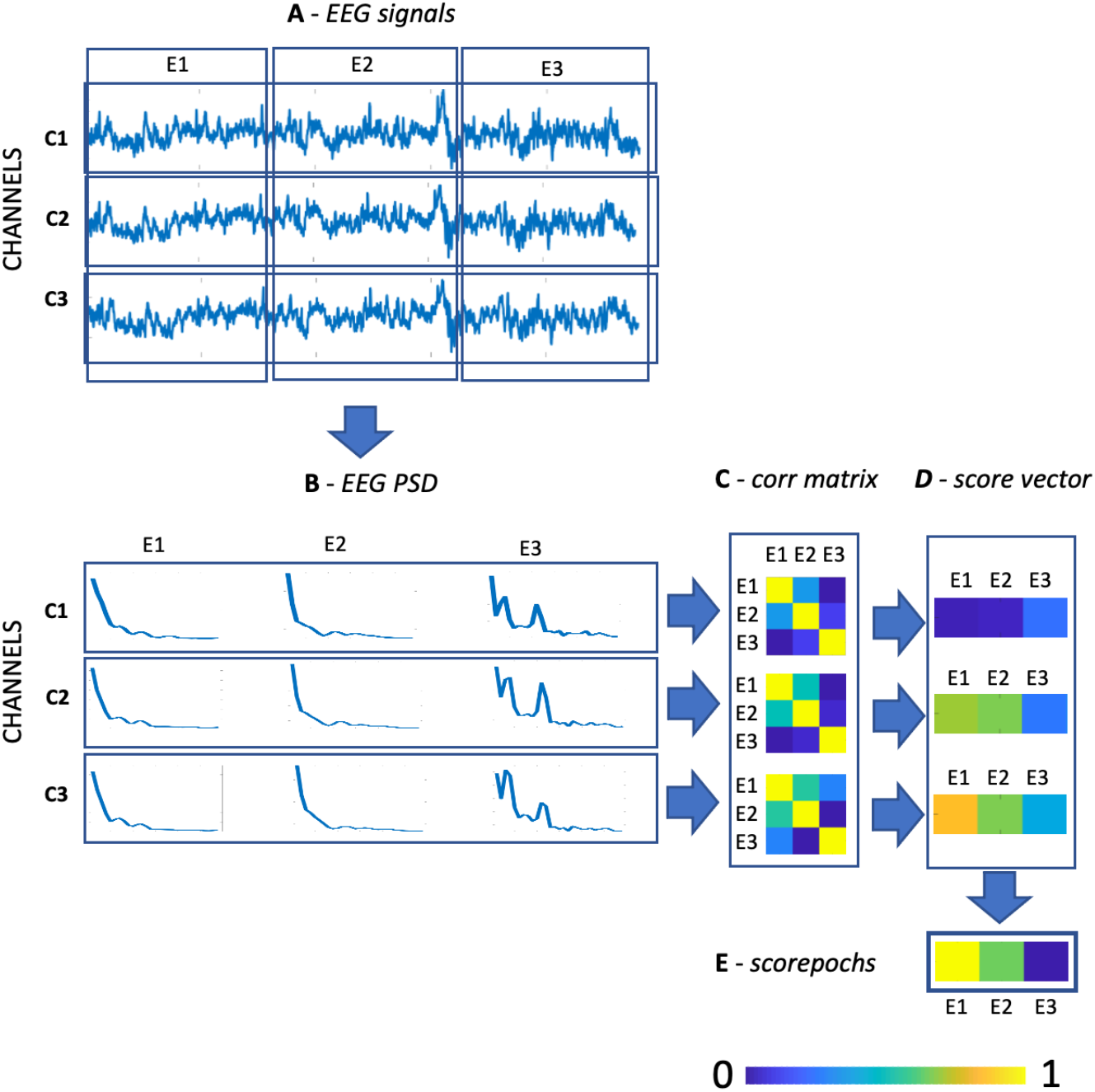
A schematic representation of the algorithm used to compute Scorepochs. Panel A: EEG raw signals of three channels with epoching scheme; Panel B: Power spectral density plots for each channel and each epoch; Panel C: correlation matrices for each channel; Panel D: score vector for each channel and for each epoch; Panel E: score for each epoch.

We tested our approach on a freely available EEG dataset [13,14] containing recordings from 109 subjects collected using the BCI2000 64-channel system (http://www.bci2000.org). The EEG dataset is available at the following link: https://physionet.org/content/eegmmidb/1.0.0/. We decided to define an easily interpretable hypothetical scenario where the interest was to contrast two different baseline conditions, namely eyes-open (EO) and eyes-closed (EC) resting-state. To contrast the two conditions the relative alpha power (computed in the range between 8 and 13 Hz) was the perfect candidate, as this property is a very common yet effective feature able to detect macroscopic differences between EO and EC conditions. The analysis was performed on 99 out of the 109 subjects since some of them were excluded due to differences in recording parameters or overall poor quality. We used an epoch length of 5 seconds and thus segmented the one-minute available recordings into twelve non-overlapping epochs (the results were successively replicated using two different epoch lengths, respectively of 2 and 8 seconds). For each epoch and each channel, we extracted the relative alpha power and the average across channels was successively evaluated for each epoch, thus computing the relative alpha power at global level. To mimic a realistic epoch selection procedure and investigate its possible effect, we decided to select for each subject four of the twelve available epochs, considering 495 different selections, representing all the possible combinations obtained by using the same subset of epochs for all the subjects. We then compared the results in terms of magnitude of effect-size obtained contrasting the two conditions (EO vs EC) on a group-level. We computed 495 t-tests, where for each test we selected 4 epochs for every subject in sequential order (i.e., for all subjects, for the first test the epochs selected were (1, 2, 3, 4), for the second test the epochs selected were (1, 2, 3, 5), for the third test the epoch selected were (1, 2, 3, 6), …, up to the last test where the selected epochs were (9, 10, 11, 12)). We then compared the magnitude of the paired Cohen’s d effect-size obtained using the selection suggested by Scorepochs against the distribution of Cohen’s d effect-size based on the sequential random selection. The analysis was later replicated using a completely different method, namely the phase lag index (PLI) [15], to compare the two experimental conditions. We performed this further analysis to investigate if the proposed approach might be successfully applied also to different methods.

Finally, to understand how much Scorepochs reflects the selection of good epochs (and not merely driven by recurring artifacts) we compared the proposed approach with another potential method based on the independent component’s classification (the number of components identified as neural is generally considered a reliable estimate of EEG signal quality [16]). For this purpose, we used the ICLabel algorithm [17] which provides - to every single component - the probability of having a cortical generator or to belong to an artifactual class (muscular, ocular, or other artifacts). Each component with a probability > 20% of having a neural source was assigned a “brain” class. The number of “brain” components was correlated - at single-subject level - with the scores averaged across all the epochs (in this case, 30 epochs of 2 second length in the 1-40Hz spectrum) as computed by Scorepochs, during the eyes-open condition. We conducted this comparison in two different and possible preprocessing scenarios. The first scenario (pipeline_01) includes the use of a bad-channel rejection approach, namely “cleanrawdata” [18], together with and artifact detection and repair method [19]. The second scenario (pipeline_02) includes the use of > 3 standard deviation approach for bad-channel rejection together with wavelength-enhanced ICA [20] for the artifact detection procedure. Later, we computed a robust statistical measure of association between the two parameters (Scorepoches versus ICLabel output) by down-weighting the outliers [21].

## 3. Results

All the results are summarized in Figure 2, which shows the comparison, in terms of Cohen’s d effect-size values, between the described sequential random selection and the selection suggested by our approach. In particular, Figure 2, Panel A, depicts the ‘effect-size time-course’ using this random selection together with the result derived by the application of our method, represented by the green dashed line. A decreasing trend in terms of effect-size values can be observed. Figure 2, Panel B, shows the distribution of the effect-size values (independently of the sequential order), where the vertical green dashed line represents the value of the effect-size obtained using the epochs suggested by our approach. The Cohen’s d value for the Scorepochs algorithm is 1.4512, which is around the 75^th^ percentile of the random epoch selection distribution. It is worth noting that the minimum effect-size is bigger than 1, meaning that, as expected, we are in a situation where the difference is largely independent of the epoch selection strategy (i.e., the difference between the two conditions EO and EC is reliably measurable).

**Figure 2.**
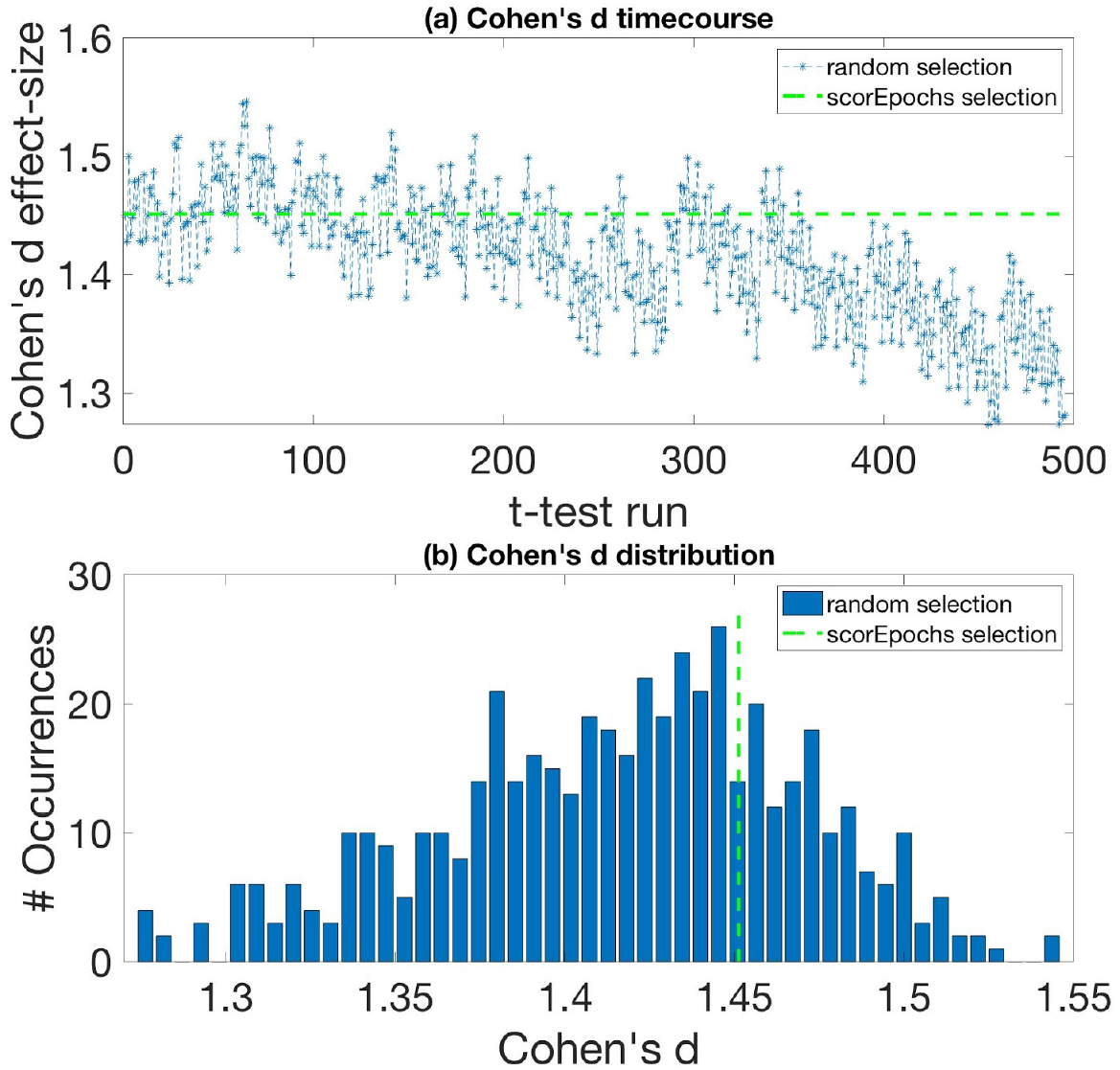
Cohen’s d effect-size for random selection and Scorepoch selection using a time window equal to 5 seconds. (a) shows the ‘time course’ of the effect-size computed using a sequential random selection. The effect-size values are reported in the y-axis, while the x-axis indicates the t-test with a sequential random selection. The green dashed line represents Cohen’s d value obtained by selecting the epochs using Scorepochs. (b) Cohen’s d effect-size distribution for random epoch selection and the Scorepoch selection. The effect-size values are reported on the x-axis, while the y-axis indicates the occurrences of specific effect-size values. The vertical green dashed line represents Cohen’s d value selecting the 4 epochs suggested by Scorepochs.

With the aim to investigate the possible effects of the time-window on the reported results, we reproduced all the analysis using two different epoch lengths of 2 and 8 seconds. The results obtained from this new analysis are represented in Figure 3 and Figure 4 respectively.

**Figure 3.**
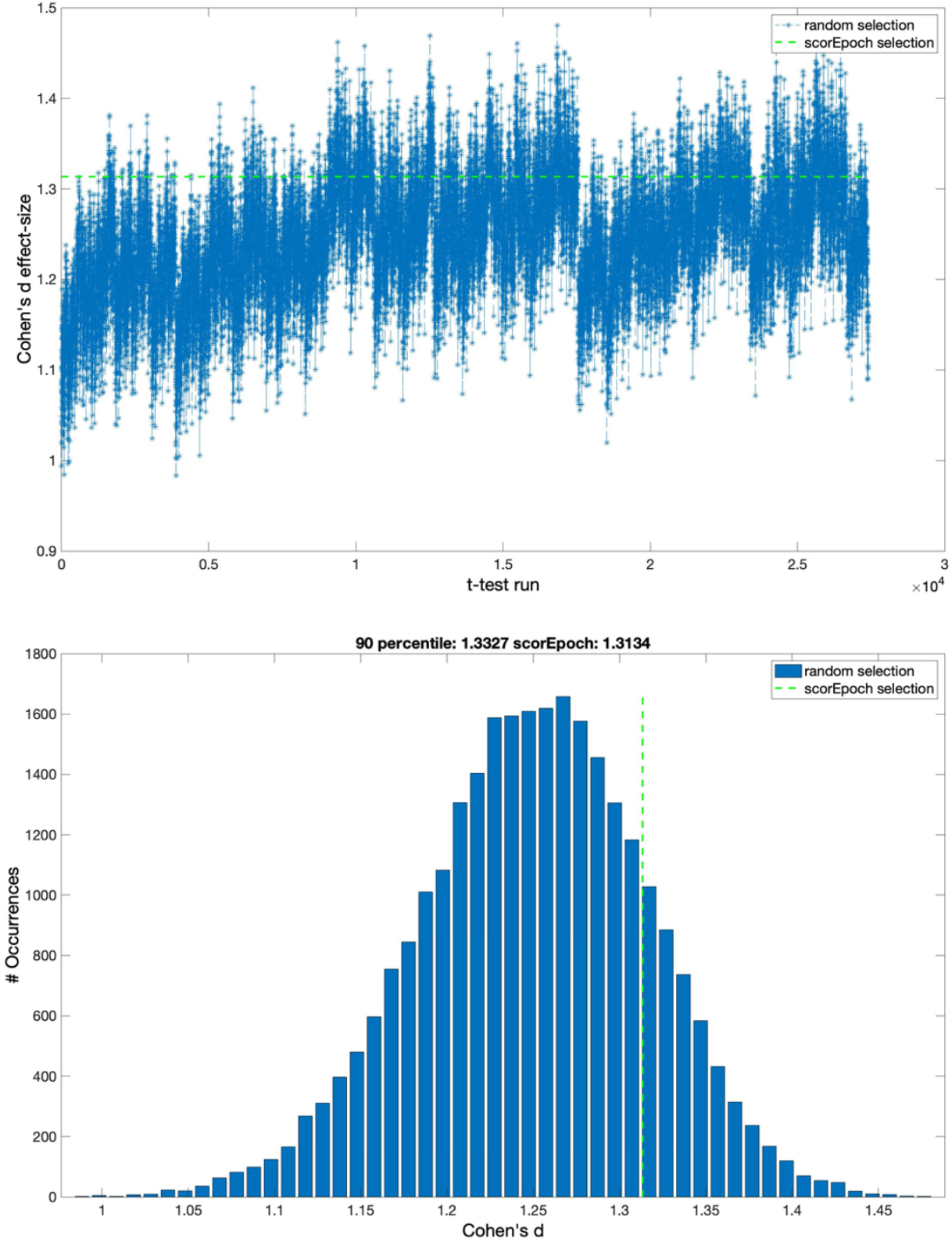
Cohen’s d effect-size for random selection and Scorepoch selection using a time window equal to 2 seconds. (a) shows the ‘time course’ of the effect-size computed using a sequential random selection. The effect-size values are reported in the y-axis, while the x-axis indicates the t-test with a sequential random selection. The green dashed line represents Cohen’s d value obtained by selecting the epochs using Scorepochs. (b) Cohen’s d effect-size distribution for random epoch selection and the Scorepoch selection. The effect-size values are reported on the x-axis, while the y-axis indicates the occurrences of specific effect-size values. The vertical green dashed line represents Cohen’s d value selecting the 4 epochs suggested by Scorepochs.

**Figure 4.**
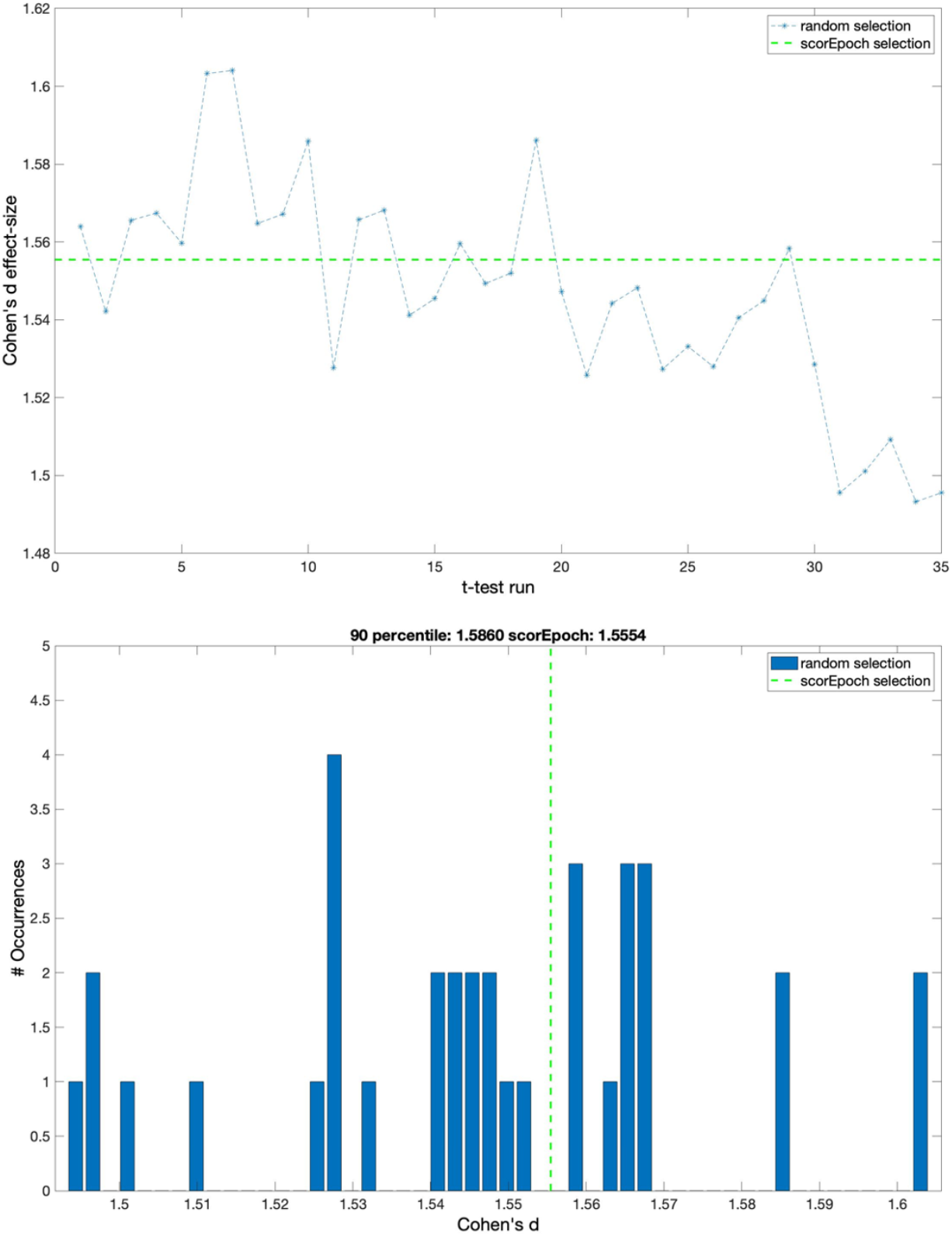
Cohen’s d effect-size for random selection and Scorepoch selection using a time window equal to 8 seconds. (a) shows the ‘time course’ of the effect-size computed using a sequential random selection. The effect-size values are reported in the y-axis, while the x-axis indicates the t-test with a sequential random selection. The green dashed line represents Cohen’s d value obtained by selecting the epochs using Scorepochs. (b) Cohen’s d effect-size distribution for random epoch selection and the Scorepoch selection. The effect-size values are reported on the x-axis, while the y-axis indicates the occurrences of specific effect-size values. The vertical green dashed line represents Cohen’s d value selecting the 4 epochs suggested by Scorepochs.

The results derived from the application of the PLI method are summarized in Figure 5 (a) and (b).

**Figure 5.**
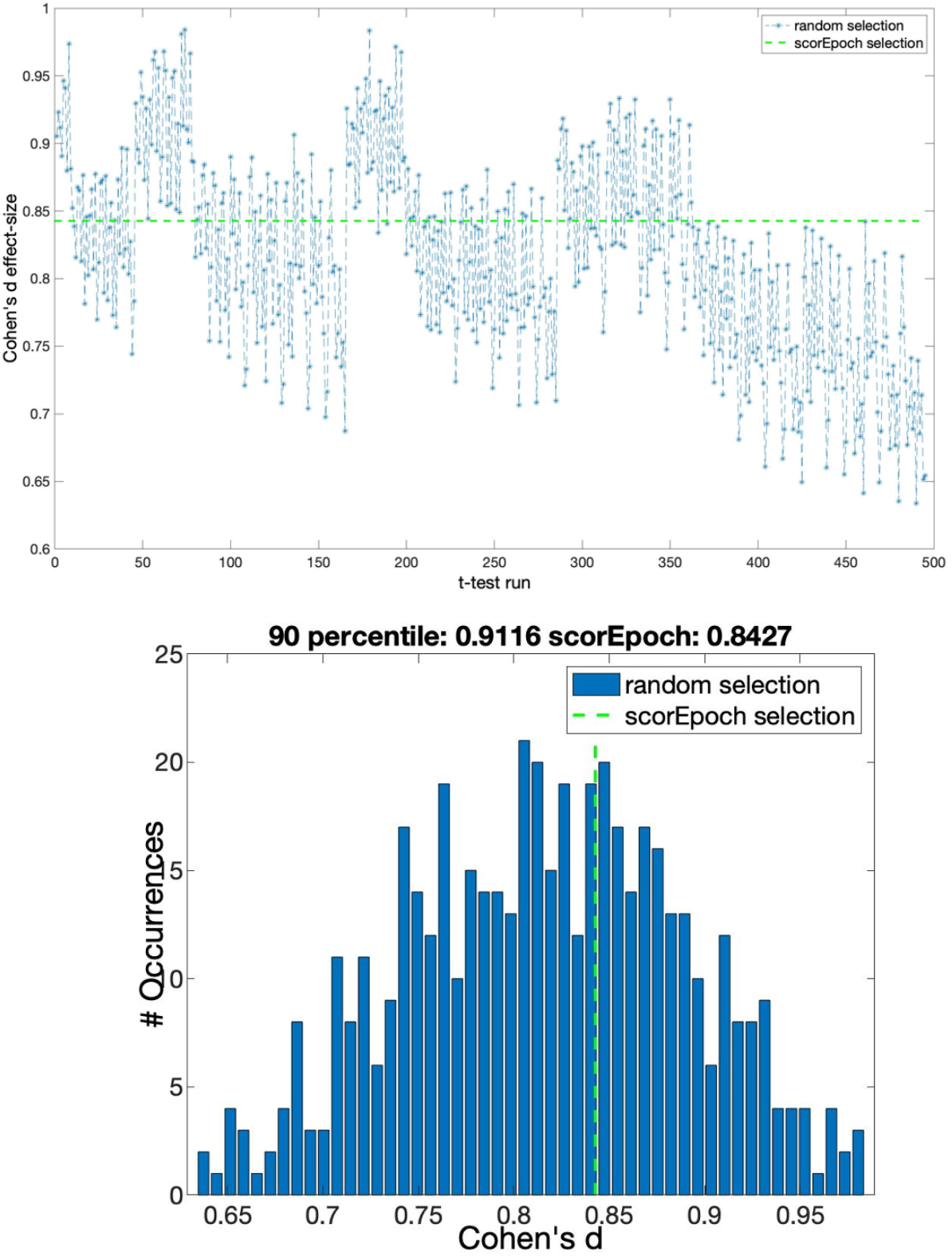
Cohen’s d effect-size for random selection and Scorepoch selection using a time window equal to 5 seconds for the PLI method. (a) shows the ‘time course’ of the effect-size computed using a sequential random selection. The effect-size values are reported in the y-axis, while the x-axis indicates the t-test with a sequential random selection. The green dashed line represents Cohen’s d value obtained by selecting the epochs using Scorepochs. (b) Cohen’s d effect-size distribution for random epoch selection and the Scorepoch selection. The effect-size values are reported on the x-axis, while the y-axis indicates the occurrences of specific effect-size values. The vertical green dashed line represents Cohen’s d value selecting the 4 epochs suggested by Scorepochs.

The last part of the analysis, as summarized in Figure 6, provides an indirect validation of the effectiveness of Scorepochs to select epochs while preserving brain activity. The comparison at single-subject level between Scorepochs and the number of independent components classified as “brain” shows a dense distribution in the upper-right portion of the scatter plots and bivariate histograms (for both the pipelines): higher values of scores correspond to a larger number of brain components. When further assessed with a robust statistical procedure such as Spearman Skipped-correlation [21], Scorepochs presents a positive association with the independent components of brain source obtained through both preprocessing pipelines (r = 0.3604, CI = [0.1780 0.5261], t = 3.9967 for the pipeline_01 and r = 0.4371, CI = [0.2856 0.5688], t = 5.0267 for pipeline_02). A similar result was reproduced when considering a stricter threshold for classifying a component as “brain” (probability > 50%).

**Figure 6.**
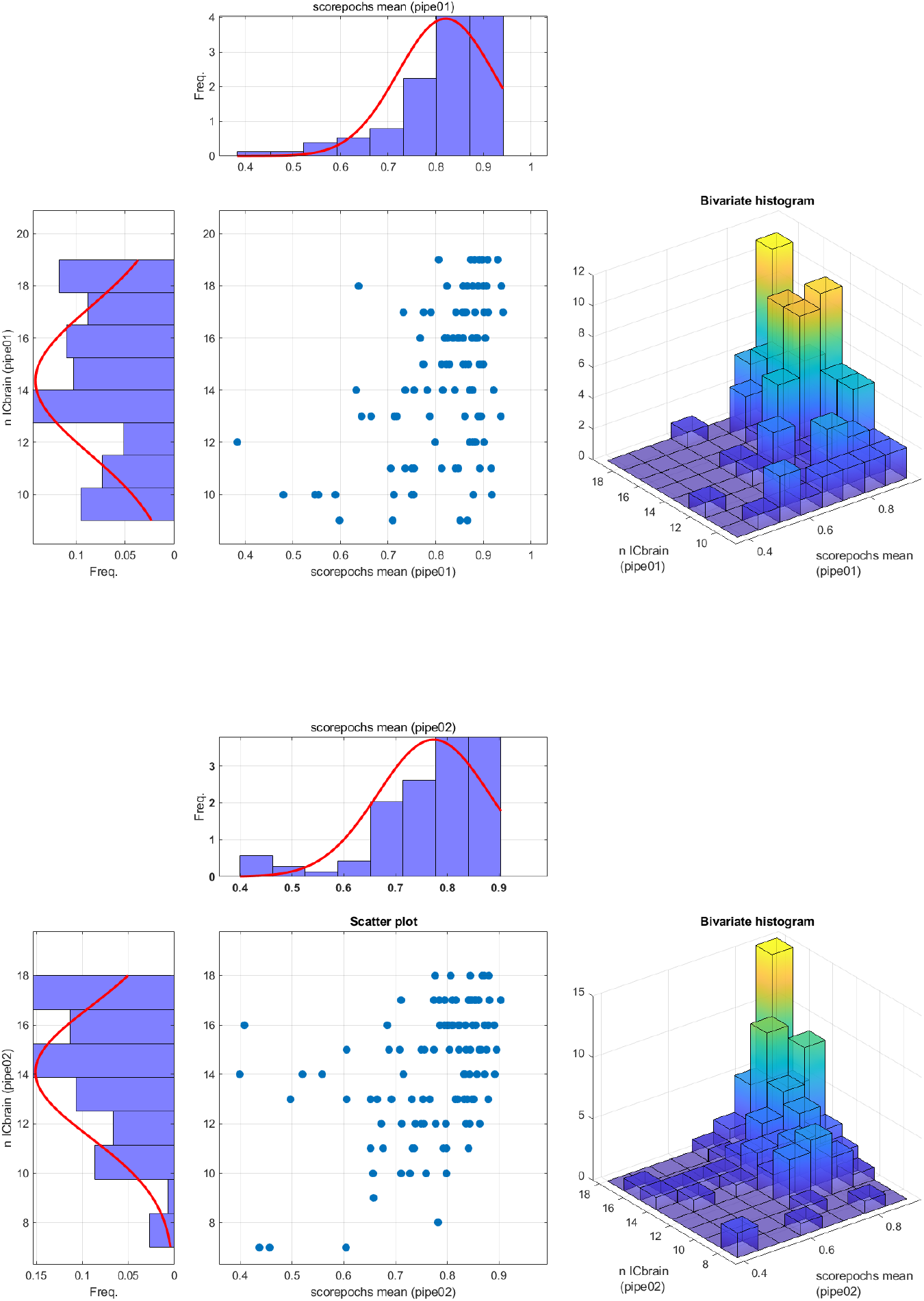
Scatterplots representing the association between the scores computed by Scorepochs and the number of “brain” components as defined by ICLabel algorithm, for pipeline_01 (upper panel) and pipeline_02 (lower panel).

## 4. Discussion

In this study, we propose an automatic method to assist M/EEG experts during the epoch selection procedure in resting-state analysis. Our method represents an objective (or less subjective) approach to perform epochs selection, if compared to the potential arbitrariness introduced by human observers and the lack of clear and shared criteria used to accomplish this crucial task. We showed, in a prototypical scenario of a group comparison between two resting-state conditions (EO vs EC), that the effect-size varied extensively depending on the epochs included in the analysis. In fact, even if it is possible to detect an effect between the two conditions EO and EC almost independently of the selected epochs (since the detection of an effect is highly probable as Cohen’s d > 1 for every epoch selection), there is a considerable variation on the effect-size depending on the actual selection. Specifically, we showed a decreasing trend (see Figure 2, Panel A) in terms of the effect-size with respect to the time in which the selection occurred (i.e., selecting epochs at the beginning of the experiment gives a higher effect-size compared to the end of the experiment). This trend may be influenced by different states of vigilance related to tiredness or drowsiness that are reflected in the recorded signals. The magnitude of the effect-size obtained using our proposed epoch selection is close to the mean effect-size (which should represent the best estimate of the population effect-size) and, more importantly, is based on a quantifiable, objective, and replicable strategy (i.e., scores computed on the PSD). It is also relevant to highlight that our results are successfully replicated using different sizes of time window. Finally, our results suggest that the proposed approach may be easily extended to other methods as the one based on connectivity metrics. Compared to other semi-automatic procedures for the selection of the ‘best artifact-free epochs’ suitable for the analysis (e.g., independent component analysis, ICA), our method is completely data-driven, and it does not require any intervention or particular skills of the user, compared to other selection strategies (e.g., knowledge of stereotypical EEG pattern related to artifact components using ICA). Moreover, the Scorepochs method, because of the small number of requirements (i.e., computation of the PSD), is not computationally expensive. Although its simplicity, this method is well grounded in physiological terms. In fact, it has been observed how the computation of simple statistics based on the PSD reflects intrinsic properties of excitatory/inhibitory levels of neuronal populations [22]. Furthermore, the PSD is able to capture different dynamics modulated by external stimuli and provides insight into sensory neural representation [23]. Finally, it has been recently reported how different behavioral states are reflected in different properties of the PSD [24]. We indirectly validated the effectiveness of Scorepochs to select good epochs by comparing our method with the number of independent components classified as “brain” using ICLabel [17], an automated electroencephalographic independent component classifier. The observed robust correlation with this approach confirms that Scorepochs may provide an objective procedure for evaluating the impact of alternative preprocessing pipelines in large-scale studies [25]. In no way, the proposed approach aims to replace the work to be performed by experts (alone or using other automatic or semi-automatic methods) during visual inspections of real M/EEG data. Scorepochs guided selection should be complementary to the human activity or to any other selection method.

## Acknowledgments

The authors have no conflict of interest to disclose. Simone Maurizio La Cava was supported by Regione Autonoma della Sardegna POR FESR Sardegna 2014-2020 Asse 1 Azione 1.2.2 CC F26C18000500006. Giuseppe Rodriguez and Matteo Fraschini were in part supported by Regione Autonoma della Sardegna research project “Algorithms and Models for Imaging Science [AMIS]”, FSC 2014-2020 - Patto per lo Sviluppo della Regione Sardegna.

